# A simple method for analyzing competitive growth of multiple cell types in xenograft tumors

**DOI:** 10.64898/2026.01.23.701386

**Authors:** Tiffany A. Melhuish, Sara J. Adair, Olivia S. Pemberton, Todd W. Bauer, David Wotton

**Author notes:** Laboratory of Molecular and Cellular Neurobiology, NIMH, Bethesda, MD, USA. Corresponding author (DW).

## Abstract

Low take rates and inter-tumor variability in growth rates can limit the effectiveness of mouse xenograft models when comparing between groups. To address this problem we developed a simple method to compare multiple cell types within a single mixed xenograft. Individual cell lines or clones were transduced with a lentiviral vector that includes a unique PCR tag, allowing the use of qPCR to determine the proportion of each tagged cell type within a mixed xenograft tumor. We generated vectors with six distinct PCR tags, and two different selectable markers, and have optimized the approach for determining their relative proportions within a mix. An initial pre-amplification step is used to increase the amount of material for subsequent qPCR reactions. This also removes the bulk of the genomic DNA, increasing the specificity of the qPCR step. Samples are then used for qPCR with specific pairs of primers that distinguish between each of the individual PCR tags, and the relative proportion of each tag is determined relative to that in the starting mix. We have tested this approach for *in vitro* growth of mixed cell cultures and in an orthotopic cecal xenograft model using a human colon cancer cell line. Since each individual tumor is initiated with a mix of cells, multiple tumors within a single animal can be analyzed separately, and overall tumor size is not important. Similarly, multiple metastatic lesions from the same animal can be analyzed individually. Thus, each tumor provides a direct comparison between individually tagged cell lines or clones. This low throughput “bar-coding” approach is simple and cost effective and has the potential to reduce the number of animals needed for xenograft experiments.

## Introduction

Xenograft models have long been an important approach for experimentally addressing the mechanisms of tumorigenesis and for testing potential therapeutic approaches in an animal model [1,2]. Human tumor derived cell lines can be implanted subcutaneously and relative growth rates measured with calipers, or tumor mass is assessed at a fixed endpoint. Subcutaneous xenografts have the advantage that they are technically easy to establish and monitor, and often have high take rates, with relatively low inter-tumor variability in growth rates. However, it may be preferable to establish orthotopic xenografts in which the tumor cells are implanted in the organ or tissue of origin, better mimicking the actual tumor micro-environment [2]. This can require more complex surgery to implant the cells, and both take rates and tumor growth rates can be much more variable. For example, orthotopic cecal tumors can be established with human colon cancer cell lines [3,4], but tumor size and end-points are quite variable [3]. This may, therefore, require larger numbers of animals to reach statistical significance when comparing between groups.

Here we present a simple approach to performing direct comparisons between genetically distinct cell types or clones, or pools of cells with different gene knock-down, knock-out or other manipulations. This approach relies on introducing two or more cell types mixed at known proportions as a single xenograft, and then determining the relative proportion of each cell type or clone in the final tumor. Thus, each tumor can be analyzed as a direct comparison between two or more cell types or clones, irrespective of how well the tumor establishes or expands. Since every individual tumor can be treated as a direct competitive growth/survival comparison, this minimizes the number of animals needed. In addition, this ameliorates problems of low tumor take rate, intra-group variability in growth rates, and variability in time to humane end-point, which may not accurately reflect tumor burden.

## Results

### Generation of PCR tag lentiviral vectors

To simplify comparison of genetically modified cell lines in xenografts, we created a series of lentiviral vectors that can be distinguished by qPCR. Cell lines with different PCR tags can be mixed at a known ratio, then after growth *in vitro* or as xenografts, the relative proportions of each cell type can be determined by qPCR in comparison to the starting mix (Fig 1A).

**Fig 1.**
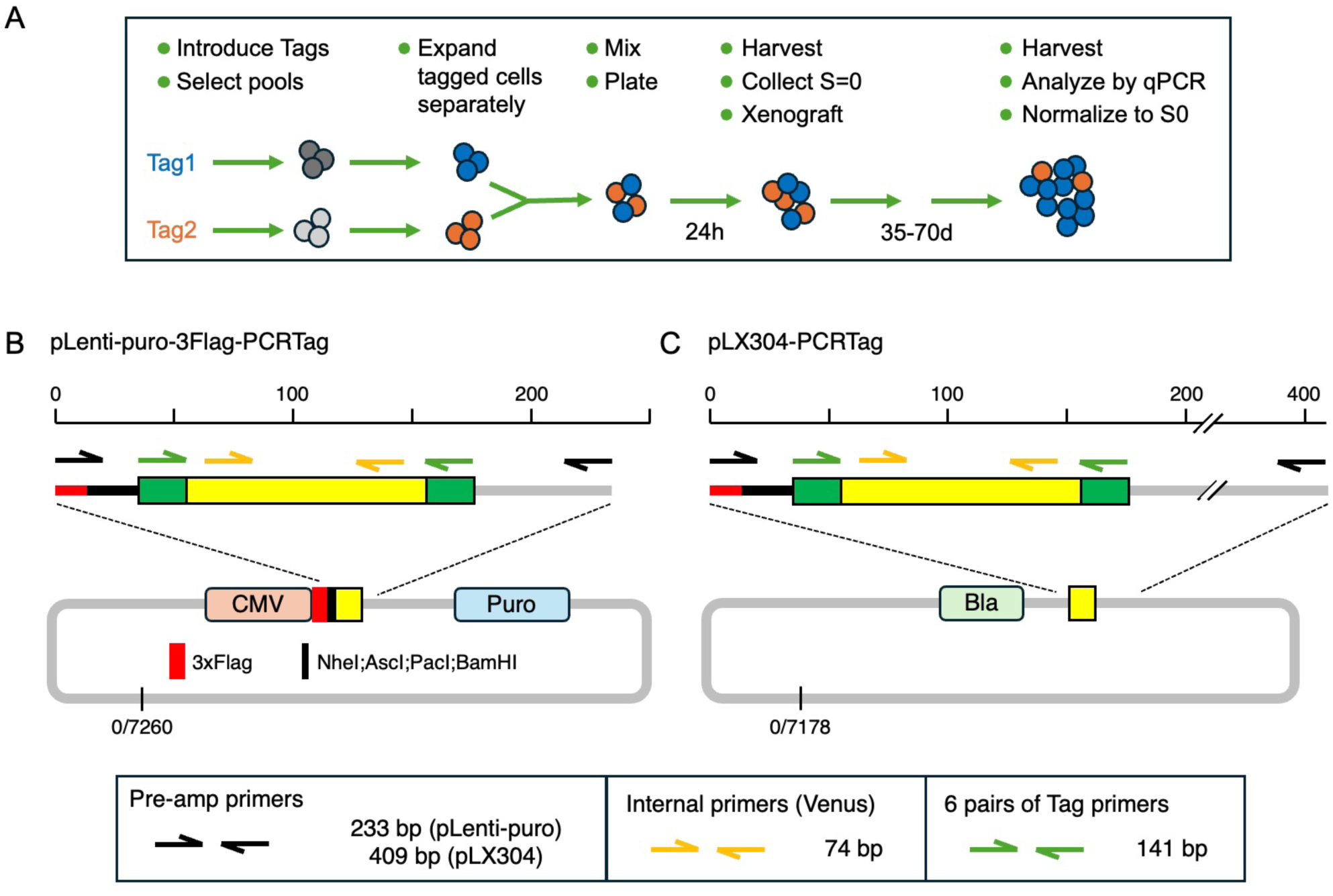
PCR tag vectors. A) the overall scheme for using PCR tagged cells in a mixed xenograft is shown. The pLenti-puro (B) and pLX304 (C) based sets of lentiviral tagging vectors are shown schematically, with the expanded regions above indicating the Tags and the locations of primer pairs.

The vectors were generated by inserting a 101bp fragment from eYFP, flanked by a pair of primer binding sites into pLenti-puro (Addgene 39481; [5]). We created a series of six different vectors with unique PCR tags, based on intron-spanning primers that we had used for qRT-PCR and had good PCR efficiency [6]. PCR with these primer pairs generates a 141bp fragment from these vectors (Fig 1B). This version of pLenti-puro also includes a triple FLAG tag and short poly-linker, allowing for gene expression independent of the PCR tag (which is not expressed). In addition, we moved all six tags into pLX304 (Addgene 25890; [7]), creating a second set of lentiviral vectors with blasticidin resistance rather than puromycin (Fig 1C).

To maximize specificity, and reduce the possibility of non-specific amplification in the qPCR step when analyzing genomic DNA samples with incorporation of the PCR tags, we designed an additional primer pair for each vector series that would amplify all six versions (Pre-amp primers, Fig 1B,C). Prior to qPCR with specific primer pairs 12 cycles of standard PCR from genomic DNA are performed and this product is purified directly on a Qiagen mini column to remove gDNA and primers. This provides enough for up to 50 qPCR reactions from a DNA sample with low complexity, and minimizes problems of non-specific amplification from genomic DNA.

### Preliminary PCR testing

To test specificity we isolated DNA from six pools of HCT116 cells transduced with one of each of the six pLenti-puro vectors, and after pre-amplification subjected each to qPCR with all six specific primer pairs and with an internal primer pair as a reference (see Fig 1). We then compared the cycle threshold (Ct) for each to the internal primer pair. As shown in Figure 2, this revealed that for each tag the specific PCR was at least 10 cycles (>1000-fold) more effective than each of the remaining five primer pairs. Additionally, by selecting specific pairs of tags this can be increased to at least 15 cycles (>30,000-fold) for comparisons of two samples.

**Fig 2.**
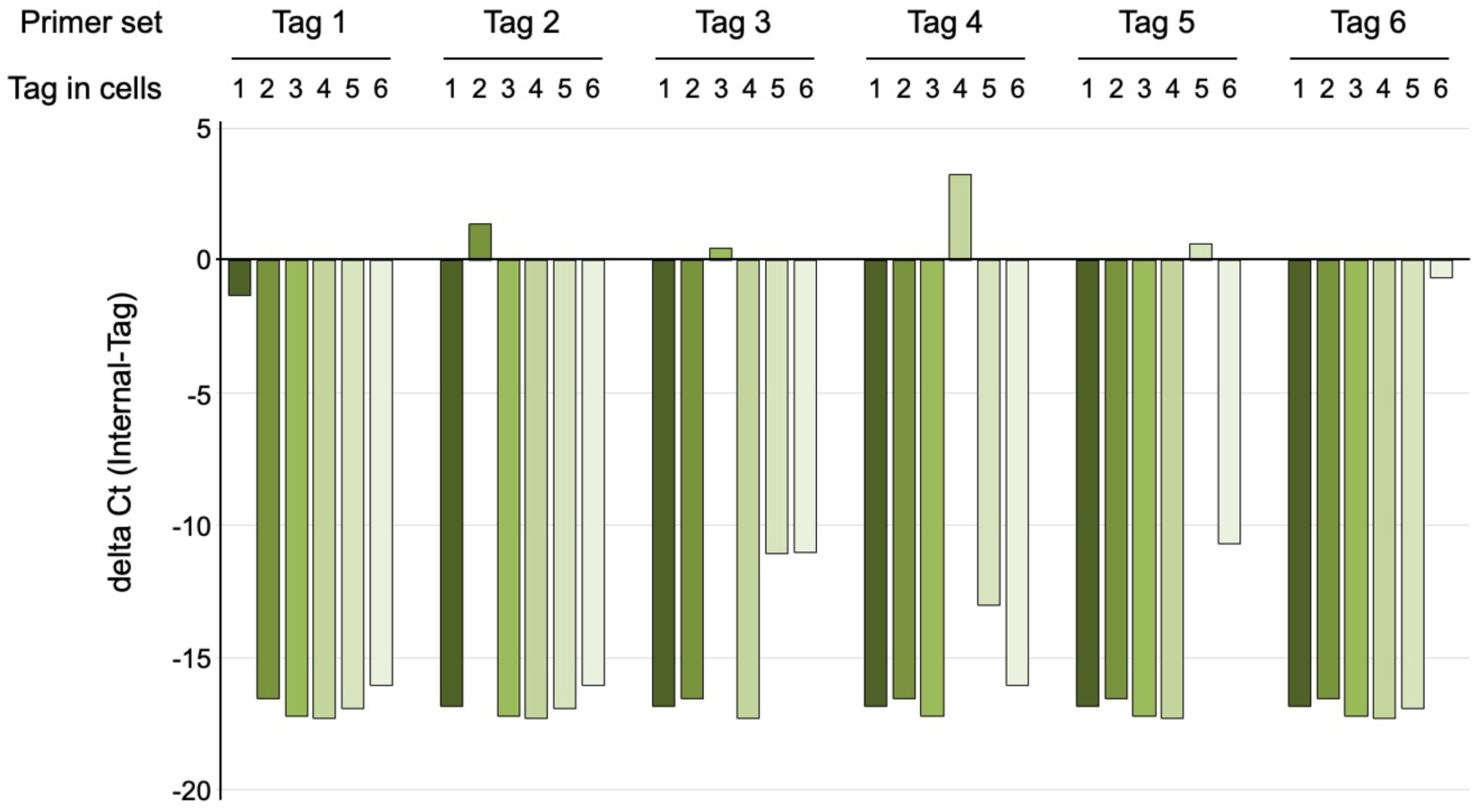
PCR tag specificity. qPCR analysis of six pools of HCT116 cells, each with one of the six pLenti-puro based Tag vectors (Tag in cells: 1-6, shades of green). Following DNA isolation and pre-amplification, each sample was analyzed by qPCR for all six Tags as well as with the internal primer pair. Data are plotted (grouped by Tag primer pair used for qPCR) as the delta Ct (internal minus Tag amplicon).

To test for linearity across a range of inputs and in different mixes, we generated a series of template mixes using a set of six lentiviral plasmids bearing each of the tags. The first mix contained equal proportions of each (∼16.7% of each PCR tag), and we also created two additional mixes, in which the amount of five of the tags remained constant while one was reduce by 100- or 1000-fold (Mix B and C; S1 Fig, panel A). Two such mixes were created for each of the six tags, and the proportion of the reduced tag in each is shown in S1 Fig (panel B, for Tag 1 as an example). The equal mix was analyzed by qPCR for all six tags and the common Venus amplicon, and each of the two reduced amount mixes was analyzed specifically for the reduced tag and Venus. These reactions were run on total DNA input amounts of 0.1ng and 0.01ng, resulting in an input of 0.00167-0.167pg of plasmid with the reduced input tag, representing 0.2% or 0.02% of total input (S1 Fig, panel B). At both input amounts, we observed linear amplification across a 1000-fold range with amplification from 0.00167pg up to 16.7pg in the equal mix, suggesting that we are able to accurately amplify a single tag form a mix in which it represents as little as 0.02% of the total (S1 Fig, panel C). To determine if the pre-amplification step affected the accuracy of the qPCR results, we performed 8, 10 or 12 cycles of pre-amplification on a genomic mix containing all six tags, then subjected each to qPCR. As shown in S1 Fig (panel D), all six tags and the common Venus amplicon were amplified linearly, suggesting that the pre-amplification step does not introduce a significant bias for or against any specific tag.

### In vitro testing

As a first test we compared all six tags together in HCT116 cells grown *in vitro*. HCT116 cells were transduced with a lentiviral shRNA vector targeting human *TGIF1* or with a control shRNA vector, and we also introduced one of the six pLX304 based PCR tags (Fig 3A). Cells were then mixed as two separate pools with equal proportions of six transduced cell types: Cells in one pool contained the control shRNA with each of the six PCR tags, and a second pool was generated in which tags 1-3 were present with shTGIF1 and tags 4-6 with the control vector. We have previously shown that Tgif1 contributes to intestinal tumor progression in a genetic mouse model [8]. In HCT116 cells *TGIF1* knockdown with this vector reduced TGIF1 protein expression by around 50% and resulted in a modest reduction in proliferation *in vitro*, with relative cell number ∼20-30% lower after five days in culture [9]. Pools were plated in triplicate (10^5^ cells/60mm plate) and passaged on days 4 and 7 to collect two sequential passages, or were plated at very low density and colonies collected on day 7 (Fig 3A). Following pre-amplification and qPCR we compared the proportions of each of the six tags to the starting mix. As shown in Figure 3B, in the pool in which all six tags were associated with the control shRNA vector, the proportions of each tag remained relatively constant in both passages and in the low-density plated cells. In contrast, the proportion of tags 1, 2 and 3 in the shTGIF1 containing mix was significantly lower than tags 1, 2 and 3 in the mix of six control shRNAs (Fig 3C, D).

**Fig 3.**
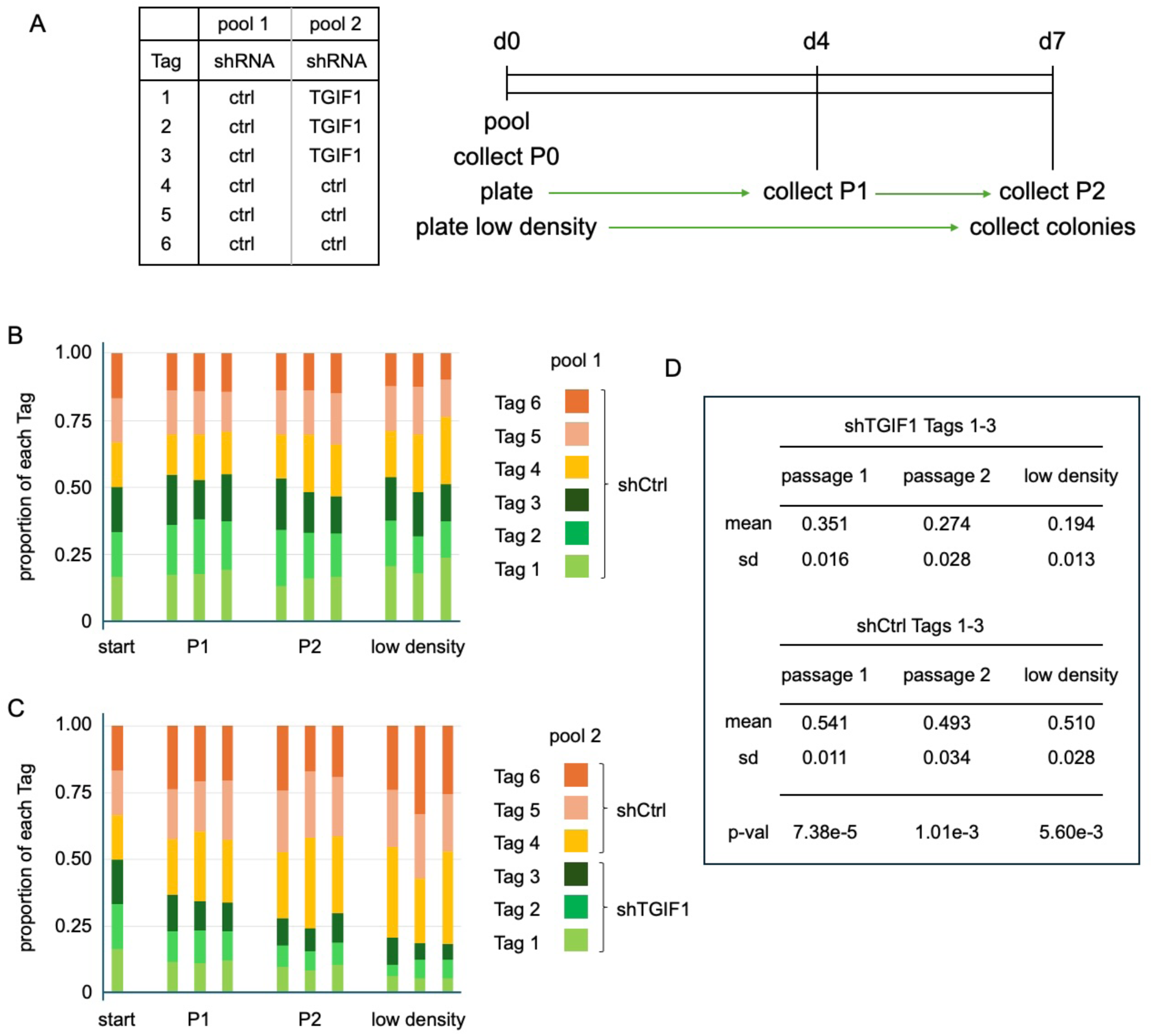
*In vitro* competitive growth analysis. A) The set-up of the two mixed pools of cells is shown – each contains an equal mix of HCT116 cells with each of the six Tags. The schematic for sample collection is shown: d0 = is the initial passage (P0). Each of three triplicate cultures was sequentially passaged on days 4 and 7 (P1 and P2). An additional set of triplicates was plated at low density and colonies harvested at day 7. The proportions of each Tag in each sample are shown for the control only shRNA mix (B) and for the shCtrl plus shTGIF1 mix (C). Note that the “start” shows the proportions of each based on the equal mix of each Tag that was used to normalize, and is shown only for comparison to the experimental. D) Comparison of the proportions of each sample that are made up of Tags 1, 2 and 3 in each of the mixes, at P1, P2 and in the low-density plated samples.

### Analysis of mixed cell type xenografts

While we can mix all six tags, the simplest approach to comparing growth is a one-to-one comparison with each cell type mixed at equal proportions, such that each sample (xenograft tumor) represents an independent competitive growth/survival assay. We set up two cohorts of 10 Foxn1 nude mice with orthotopic xenografts, in which a total of 10^6^ cells were injected into the wall of the cecum divided between six separate sites. Each mix contained equal proportions of shControl and shTGIF1 cells, either with tags 5 and 2 (Fig 4A) or 6 and 1 respectively (Fig 4B). One cohort was allowed to reach humane end-point (or a maximum of 70 days, Fig 4B), and the other was collected at 42 days (Fig 4A). From each group, 9 mice survived and had identifiable tumors when euthanized. In the first group we found 16 tumors at 42 days, since some mice had clearly separable tumors resulting from the multiple injection sites. In all of these, the control cells had outgrown the shTGIF1 cells (Fig 4A). In the second group that were analyzed at a later time, we found 13 clearly separable tumors, and all but two had a majority of control cells (Fig 4B). Interestingly, the one tumor in which shTGIF1 cells made up more than 90% was the smallest of all the tumors isolated. We found similar results, with selection against shTGIF1 cells in pools in which we used all six tags, with three each for shTGIF1 and shControl [9]. Together these results suggest that similar results are obtained independent of which PCR tags are used, and that the approach is quite reproducible.

**Fig 4.**
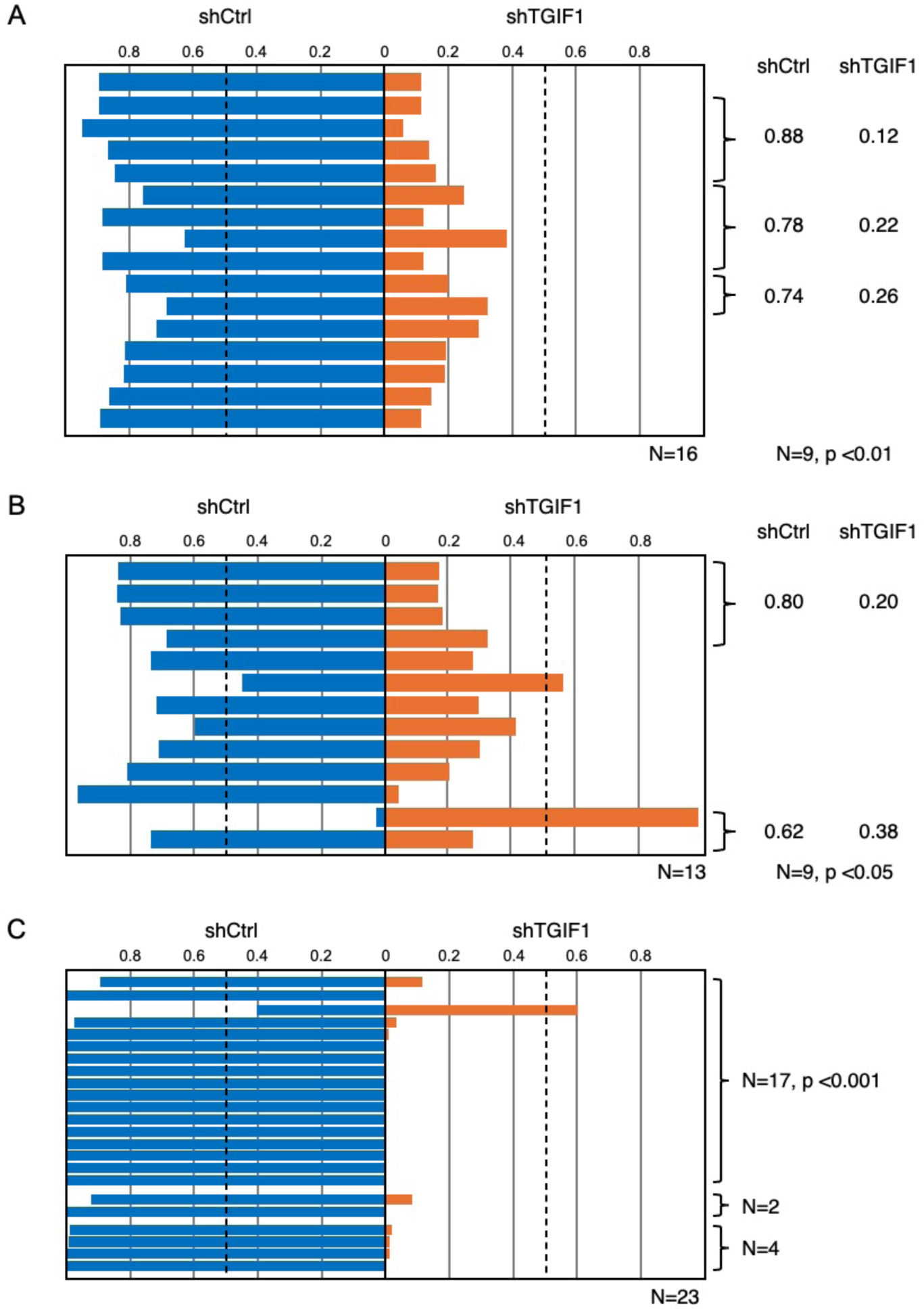
Analysis of orthotopic xenografts. Orthotopic cecal xenografts were generated using an equal mix of each of two cell pools, one with a control shRNA, the other with shTGIF1. Each also contained a distinct PCR tag. The proportion of each Tag (associated either with shCtrl or shTGIF1) in each separate tumor are plotted for two cohorts of mice. A) Mice were euthanized at a maximum of 6 weeks after xenograft (16 tumors from 9 mice). B) Mice were allowed to reach humane end-point or a maximum of 10 weeks (13 tumors from 9 mice). C) Individual liver metastases from the two cohorts of mice in A and B, were analyzed similarly. Tumor number is shown below each panel, and the proportion of each tag averaged per mouse, for mice with multiple tumors (indicated by brackets) is shown to the right (A and B). The p value (Wilcoxon signed rank, two tailed) is shown (analyzed by animal, N=9). In panel C, the p value (Wilcoxon signed rank, two tailed) for the 17 metastases from one of the mice is shown.

From the two cohorts of mice analyzed here we also identified 23 spontaneous liver metastases, and all but one of these had a preponderance of shControl cells (Fig 4C). From the shTGIF1/shControl tumors in which all six tags were present, we were able to test for the presence of each of the six independent cell lines in individual tumors and metastases. As described here for the one-to-one comparisons (Fig 4), shControl cells were the majority in all but two of 25 metastases found in a separate cohort of mice in which all six tags were present [9]. The analysis shown in Figure 5 suggests that these cells metastasize not as single cells but rather as multi-cellular clusters, and suggests that including multiple tags allows for the possibility of following multiple cell types in a single tumor and in the resulting metastases. For example, a mouse in which two primary tumors were present, with all six tags represented between the two gave rise to metastases with exclusively control cells, but that contained a mix or two or three of the different control lines (Fig 5A). The second mouse shown here had a single primary tumor with less than 5% shTGIF1 cells, but gave rise to metastases that were predominantly either control or ∼90% shTGIF1, in one case (Fig 5B). Similarly, metastases with various combinations of five of the six tags were identified from a mouse in which the primary tumor retained all six tags (Fig 5C). The three mice analyzed here were selected to show situations where the final mix in specific metastases did not necessarily match that in the final primary tumor, and where multiple tags were present. This analysis suggests that combining multiple tags retains the ability to perform one-to-one comparisons as with the use of only two tags, but also provides for the possibility of comparing multiple groups and giving some insight into the complexity of the tumor.

**Fig 5.**
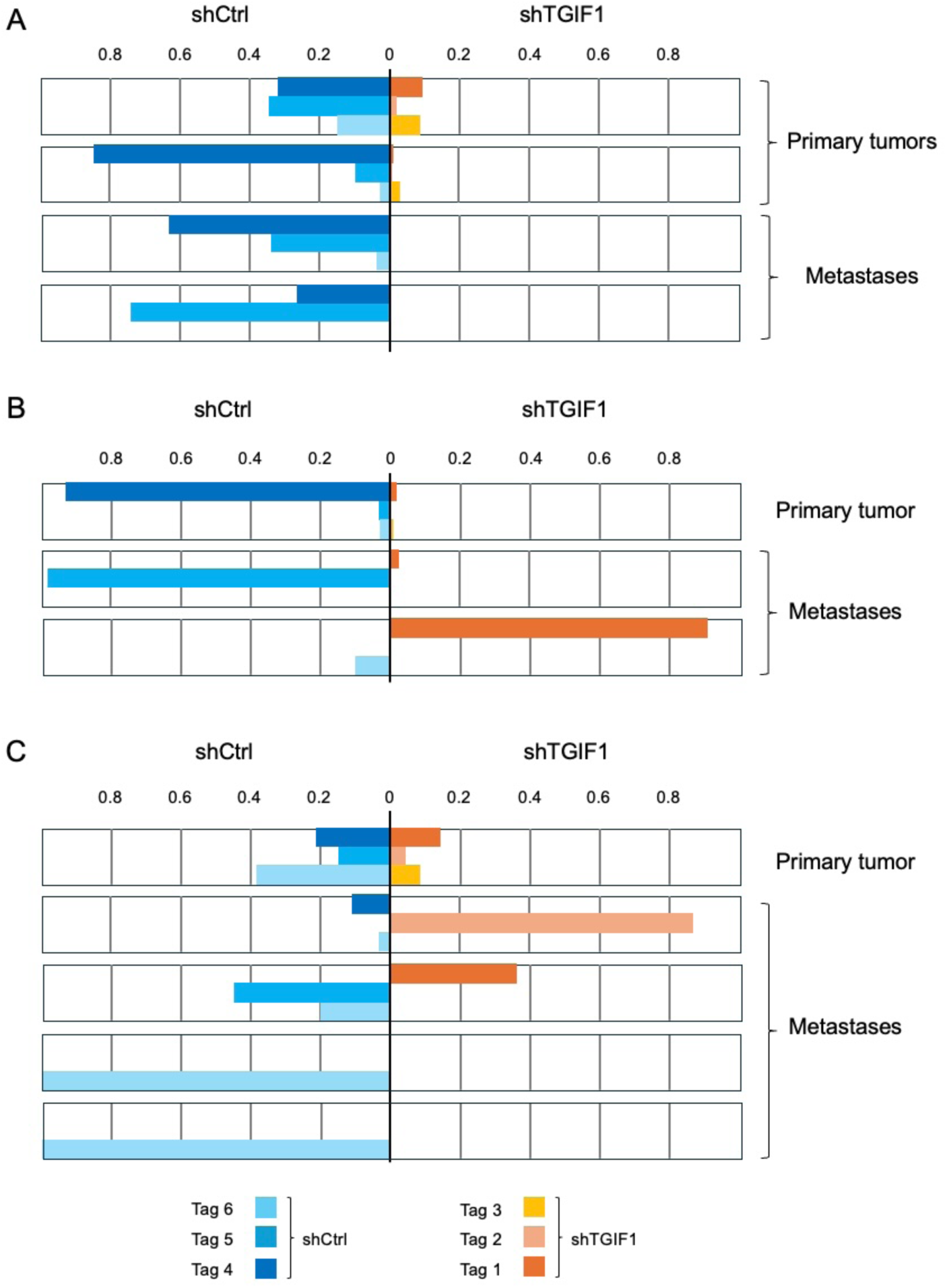
Analysis of tumors with multiple Tags. Data from three mice is shown, in which a mix of six cell pools was used, with all six Tags: Tags 1-3 with shTGIF1 and Tags 4-6 with shCtrl. In each case the proportion of each Tag in individual tumors and liver metastases is shown with primary tumors above and metastases from the same mouse below. The mouse in presented A had two separate primary tumors with different amounts of shTGIF1 cells, and two liver metastases without any shTGIF1. Mouse B had a single primary tumor with very little shTGIF1 at end-point and two very different liver metastases. Mouse C had a single relatively mixed primary tumor with four liver metastases of differing make-up. Note this analysis is of a xenograft experiment described in reference 9.

### Statistical analysis of mixed xenografts

The xenograft data with shTGIF1 described in Figure 4 was compared using a Wilcoxon signed rank test, in which we averaged the proportion of tags present in primary tumors for each animal to avoid problems with pseudoreplicates. In both cases, there was a significant enrichment of shControl cells when analyzing the data with multiple tumors averaged by animal (p < 0.01 and p < 0.05; two-tailed Wilcoxon signed rank). For some xenograft approaches, only a single tumor per animal will be generated so dealing with pseudoreplicates will not be an issue. If only a small fraction of animals have multiple tumors, then averaging tumor data (or pooling tumors) by animal may be reasonable, as in Figure 4 A and B. For analysis of the metastases (Fig 4C), only three mice had spontaneous metastases, such that averaging per animal does not provide a large enough N for the Wilcoxon signed rank test. One option here is to analyze data from each mouse separately, but this also reduces the numbers: In the case shown in Figure 4C, only one animal had enough metastases for analysis by Wilcoxon signed rank.

Where multiple tumors or metastases are found in most animals, and there is a desire to reduce animal numbers, we suggest using the internal Venus amplicon (which is present in all tags) in addition to using the tag-specific primers, then performing a two-way ANOVA with the animal as a covariate. This allows for the analysis of multiple tumors isolated from smaller numbers of animals. The ΔCt (starting mix – tumor, for a specific tag) is normalized to the Ct value for the Venus primers for each sample to account for differences in input and the ANOVA test is run on this ΔΔCt data. As an example of this kind of analysis, we compared data from nine cecal tumors obtained from four mice, injected with a mix of two pools of HT29 colon cancer cells (S1 Table) infected with either Tag 3 or Tag 6 and shControl or shACADM lentiviruses. While the data was significant with a simple Wilcoxon signed rank test (p = 0.007812), this does not take into account pseudoreplicates, and because these tumors came from only four mice, this is too few for meaningful comparison by Wilcoxon signed rank if averaging tumors by animal. However, when we compare the nine tumors by multiway ANOVA with the animal identity as a covariate the data are clearly significant (p = 0.000152, by tag; S1 Table). This approach allows for statistical comparison of multiple tumors or metastases from small numbers of animals that would otherwise not be comparable.

### Optimized method for analyzing PCR tags

Based on our experience comparing different approaches to using these PCR tags, we have come up with a relatively simple optimized approach (Fig 6). The initial set up involves expanding the cells separately, then mixing and plating for overnight growth prior to harvesting for experimental set up. This ensures that all cells have been treated equally on the morning of setting up xenografts, as they are trypsinized and collected as a single pool. In addition to collecting a “time 0” sample immediately prior to generating xenografts (Fig 6A), we generally also collect one at day -1, when plating the initial mixed cell pool, and from the cells and at day +1, by plating a fraction of the cells left over after generating the xenografts. In our experience all three of these (days -1, 0 and +1) have been very similar to each other, even in cases such as the shTGIF1 versus shControl, where shTGIF1 cells have some growth disadvantage *in vitro*. However, comparing these three control samples ensures that no problems have arisen between the initial mixing and the end of implanting the cells.

**Fig 6.**
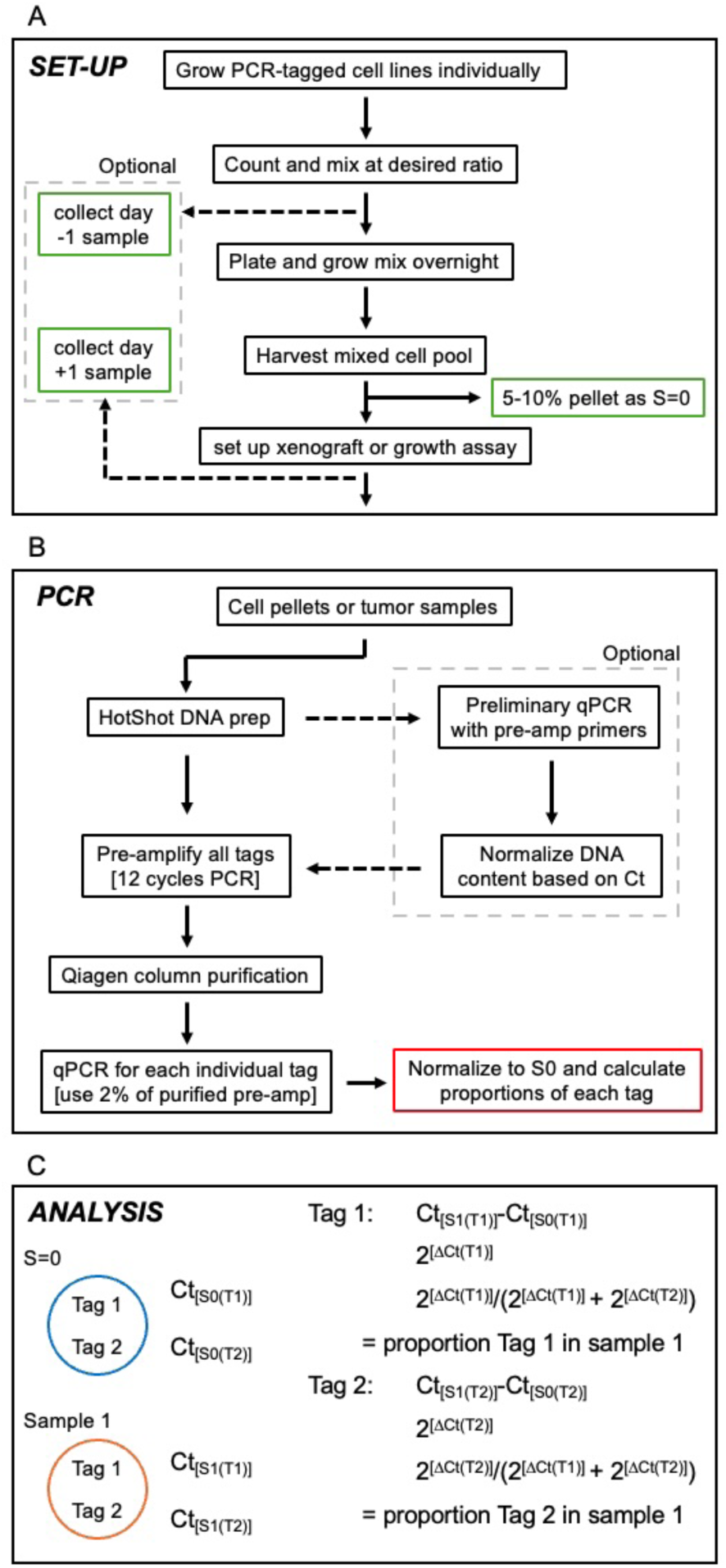
Optimized approach. The optimized approach for using these PCR tags to compare growth and/or survival of different cell types in mixed xenografts is shown schematically (see also Materials and Methods for a detailed description). A) the set-up for generating the initial mix and collecting sample 0 is shown. Optional collections at one day before and one day after xenograft can be useful to ensure all cells are well represented and heathy at the start. B) The scheme for pre-amplification followed by qPCR is shown with an option to first normalize samples by qPCR with the pre-amp primers. C) The ΔCt method to calculate relative proportions in each sample is shown.

Once all samples are collected, we perform a quick and simple DNA prep by HotShot [10], which simply involves boiling the sample in sodium hydroxide, then neutralizing with Tris. This is adequate for direct use in the pre-amplification PCR. Optionally, for samples that are very different in size, or where there are concerns of different amounts on contamination by non-tumor tissue, an initial qPCR run can be performed (with either pre-amp or internal primers) and used to normalize the amount of HotShot sample to be used for the 12 cycle pre-amplification (Fig 6B). Following Qiagen spin column purification of the Pre-amp PCR, we then use 2% of the eluate for each qPCR reaction. We have found that there is no need to test how well the pre-amplification and purification has worked. Occasionally, for samples such as extremely small metastases we get no product, but this is relatively rare, and some variability in the amount of input for the qPCR can be tolerated.

Analysis of the qPCR data uses a ΔCt method to calculate a proportion of each tag in each experimental sample, relative to that in the starting sample (S=0) (Fig 6C). The proportions at the start (S=0) are assumed to be at the desired ratio based on the initial mixing of the cells (50% each for 1+1, 16.67% each for all six tags at equal proportions, for example). Including an analysis of the day -1, day 0 and day +1 at this point provides a good comparison of samples that are expected to be very similar, and ensures that no problems occurred during the setup. For statistical comparisons between two groups we have used the Wilcoxon signed rank test for comparing the proportions in xenografts. This requires around 8-12 tumors – eight is enough to give statistical significance if the majority are of one type. As for the cell line data shown in Figure 3, it seems reasonable to compare the proportion of the sample with the test cells to the proportion of the same tag in a mix of all control cells using a student’s T test. This seems less appealing for xenografts, since it would require setting up a second cohort in which all cells are control cells with at least 2 different tags. Within a mix of six tags, such as the 3 + 3 used in Figure 3 it is probably not appropriate to compare the three experimental tags to the three controls in one tumor since if the experimental tags decrease, then the control have to increase as a proportion.

A more detailed description is provided in the materials and methods section, together with details of the vectors and primers used.

## Discussion

Here we describe a relatively simple PCR based method for comparing competitive growth/survival in a mixture of different cell types either *in vitro* or *in vivo*. Conceptually, this is based on the widely used bar-coding approach in which thousands of distinct samples are analyzed together and distinguished by high-throughput sequencing. Our method is relatively inexpensive and is suitable for comparisons of small numbers of samples in a mix, for which library generation and sequencing would be cost-prohibitive. This approach is ideal for comparing two genetically modified or otherwise distinct cell types. By reducing the numbers of mice needed, this may be particularly helpful in xenograft approaches where tumor take rates are low, growth rates are variable, or the generation of orthotopic xenografts is technically challenging and time-consuming. If multiple tumors can be isolated from a single animal, this has the potential to further reduce animal numbers, since each can be analyzed separately, rather than simply measuring overall tumor burden or time to end-point for the animal. Additionally, this approach has the potential to reveal relatively subtle differences in tumor growth that might otherwise require much larger numbers of animals. While this approach may be advantageous for comparison of tumor growth rates, it is worth pointing out that since most resulting tumors will contain a mix of the starting cell types, they are unlikely to be useful for the analysis of gene and protein expression, or for histological analyses. However, this approach provides a powerful way to generate tumor growth data from a small number of mice, and additional unmixed tumors can be generated for other analyses.

We show that this method appears to be linear across a wide range of proportions of each tag, down to 0.02% of the total tags present. In a situation in which two tags are initially mixed at equal proportions, this allows for detecting a reduction of ∼2500-fold. Using the pre-amplification step reduces the possibility of non-specific amplification from genomic DNA in the qPCR step. Importantly, the pre-amplification step does not appear to introduce any significant bias and increases sample availability for qPCR from relatively small initial samples. The pre-amplification step, followed by column purification also allows us to use a relatively quick and simple initial DNA isolation procedure, and avoids potential concerns with the relative amounts of tumor cells to non-tumor cells in the starting samples.

For the analyses described here we have used pools of cells as soon after lentiviral infection as possible, to avoid potential confounding effects of prolonged time in culture to isolate individual clones and possible concerns with clone-to-clone variation. However, isolating multiple clones and comparing results among them may also be a good approach. In this case, it might be appropriate to use all six tags, which would allow for comparing three control and three experimental clones in a single mix. One possible concern with this is that if there are significant differences between clones, then a single clone might take over at the expense of others and make interpretation more difficult. Perhaps a better approach would be to use three tags each for control and experimental, but to combine them pairwise as a series of one-to-one tests. Other scenarios in which we could envision using multiple tags would be as an initial screen of a small number of candidates, for example, comparing five experimental cell lines with one control as a first test of whether any of the five perform better or worse than the control. As with the analysis in Figure 5, the use of multiple tags within a single mix can also be used to examine how multiple cell types behave in a more complex situation rather than simply comparing growth or survival at a single site.

For the analyses shown here we used two vectors, the puromycin resistant shRNA vector and the blasticidin resistant pLX304 Tag vector. For expressing a gene of interest and introducing a PCR tag, there are two options. First a two vector set-up, with either the pLX304 or pLenti-puro carrying the PCR tag, depending on compatibility with the expression system. The second option is a single plasmid approach, in which the gene of interest is expressed from the pLenti-puro Tag vector, either using the included triple Flag tag, or after removing it. However, in this case an insert specific pre-amplification primer would be needed, and this should be common to all inserts in a comparison. Additionally, it should be noted that in our experience expression from pLenti-puro is relatively low, generally within 2-fold of the endogenous protein, compared to the 5- to 10-fold overexpression we have seen with pLX304. One additional approach we have begun to use is to have genes synthesized for insertion into a lentiviral vector and to include the 233bp pre-amplicon region from pLenti-puro downstream of the expression cassette within the synthesized gene fragment. This allows for gene expression or knock-down together with the introduction of a specific PCR-tag using a single lentiviral transduction.

In summary, we describe a relatively simple method for comparing growth of genetically distinct cell types as xenografts, that is particularly useful where tumor growth or the humane end-points are variable, or where the tumors do not become large enough for accurate measurements. In such cases, this approach should simplify comparisons and minimize the number of mice needed.

## Materials and methods

### Detailed method for use of PCR tags

#### A) HotShot DNA preps

##### Cultured cells

Cell pellets collected by scraping or trypsinization were resuspended in PBS at 10^7^ cells per ml. Pellets can be stored at -20°C prior to resuspending. Genomic DNA was prepared by HotSHOT [10], diluting resuspended cells 1:6 (20μl + 100μl) in HotSHOT lysis solution (25mM NaOH, 0.2mM EDTA, pH 12), boiling (40 minutes) and neutralizing with equal volume HotSHOT neutralization solution (40mM Tris-HCl, pH 5).

##### Primary tumors

For larger samples, such as primary tumors, they were collected, weighed, frozen on dry ice and stored at -20°C. Cecal tumors were homogenized using a Polytron homogenizer at 300 mg/ml in PBS on ice. Genomic DNA was prepared by HotSHOT, diluting homogenized tumor 1:6 in HotSHOT lysis solution, as for cell pellets. We have also used a similar approach for subcutaneous xenografts. If the primary tumors are very small they could be processed as for small metastases (below).

##### Metastases

Smaller samples such as individual metastases were resuspended in ∼8 volumes of HotSHOT lysis, boiled for 40 minutes and neutralized with equal volume HotSHOT neutralization solution. Pre-amplification from metastases of at least 1mm in diameter is routinely successful. Larger metastases can be weighed and treated as for the primary tumors.

#### B) Optional normalization

This optional normalization is not absolutely necessary, but may be useful with larger tumors where differing amounts of the sample may be non-tumor tissue. For cell pellets where approximately equal numbers of cells are present it can be omitted. Additionally, for smaller samples such as individual metastases (< ∼2mm in diameter) we omit this step, and use the HotSHOT sample directly for pre-amplification, without normalization.

<u>Pre-amp qPCR normalization:</u>

1μl HotSHOT sample

1μl 5μM each forward + reverse Pre-amp primers (200nM final of each primer)

12.5μl 2x SensiMix SYBR + FITC (Bioline)

Water to 25μl total volume

<u>qPCR program:</u>

8 minutes, 30 seconds, 95°C

40 cycles:

10 seconds, 95°C

10 seconds, 60°C

10 seconds, 72°C

Samples were then normalized around an arbitrary Ct of 30 (this can be adjusted if all or most samples cluster well below or above this). A Ct of 30 (in this example) would then require 1μl of HotSHOT for the subsequent pre-amplification reaction, with other samples scaled accordingly. The Ct of 30 is designed such that the pre-amplification reaction will bring this into the low to mid 20s range for the Ct for a tag that constitutes half the sample: 12 cycles of pre-amplification, and using only 1/50^th^ of the eluted sample for each qPCR, with each tag starting at 50% in an equal 1 + 1 mix. It should also be noted that this normalization does not need to be very accurate, since each sample is internally controlled and a relative proportion determined. This just ensures that most qPCR reactions generate a Ct that is in the optimal range for observing differences.

#### C) Pre-amplification

The PCR tag region is enriched by 12 cycles of pre-amplification PCR with primers common to all six PCR tagging vectors using Mango Taq DNA polymerase (Bioline). Pre-amplified PCR product is purified using Qiaquick purification kit (Qiagen #28104) and eluted in 50ul. This eluate can be used directly for qPCR, without further purification or first determining the concentration.

<u>Pre-Amplification PCR:</u>

1μl HotSHOT sample

4μl 5x Bioline Mango Buffer

0.6μl MgCl_2_ (50mM)

0.4μl dNTP mix (containing 10mM each dNTP)

0.2μl forward primer (100μM)

0.2μl reverse primer (100μM)

0.2μl Bioline Mango Taq

Water to 20μl total volume

<u>PCR program:</u>

3 minutes, 94°C

12 cycles:

30 seconds, 94°C

30 seconds, 60°C

30 seconds, 72°C

3 minutes, 72°C

<u>Pre-amp purification:</u>

Add 5 volumes Qiagen buffer PB to pre-amp PCR product

Apply to Qiaquick column

Wash with 750μl Qiagen buffer PE

Add 50μl Qiagen buffer EB, warmed to 60°C, incubate 5 minutes

Spin to elute purified, pre-amplified DNA

#### D) Quantitative PCR

1/50 of the total pre-amplified PCR product per well is used as input for qPCR using SensiMix SYBR + FITC PCR Mix (Bioline) and myIQ thermocycler (Bio-Rad). In general at least duplicate wells are preferred, with separate reactions for each tag in the mixed sample. Ct values are determined by IQ5 software (Bio-Rad).

<u>Individual tag qPCR:</u>

1μl HotSHOT sample

1μl 5μM each forward + reverse Tag primers (200nM final each primer)

12.5μl 2x SensiMix SYBR + FITC (Bioline)

Water to 25μl total volume

<u>qPCR program:</u>

8 minutes, 30 seconds, 95°C

40 cycles:

10 seconds, 95°C

10 seconds, 60°C

10 seconds, 72°C

#### E) Calculation of relative proportions

The proportions of each PCR tag present in the final cell pellets or tumors are calculated by 2^ΔCt^ using the sample zero (S=0) mixture of cells as the reference. For example, for two tags and a single sample compared to a starting mix:

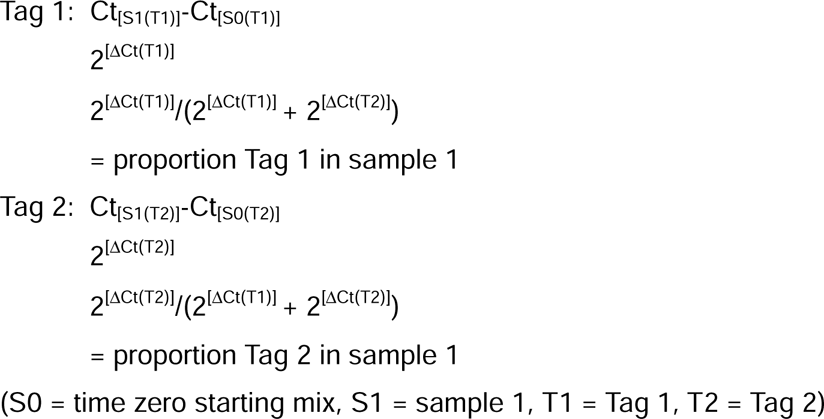

#### F) Primers and vectors

qPCR tag primers:

PCR tag 1 F: ACAAGTGCGGAGTATGTGGA, R: TCTTTGGCTTTGAACTGTCG

PCR tag 2 F: TACCAATGACAACGCCTCCT, R: TGGGAGTACGGATGCACTTT

PCR tag 3 F: TGAATGGACACTCTGGGTTG, R: CGATCAGTGGAGGTCTTTGG

PCR tag 4 F: TGAACCAGGAGATCCGAGAC, R: GAAACAGGGCTTCATTCTGC

PCR tag 5 F: CCAGATGGAAGCCGATATGA, R: TCGTTCTGCTGCTCAATCAC

PCR tag 6 F: TCCCTCTGTGATGAGCTGTG, R: TCCTCTTCCTCGGTGTCTTC

Pre-amp primers:

pLenti-puro F: TGATGATGATAAGGCTAGCGG, R: CCGAGGAGAGGGTTAGGGAT

pLX304 F: GGGACAGCAGAGATCCACTTT, R: GGGCCACAACTCCTCATAAA

Internal primers:

Venus F: CAACTACAACAGCCACAACG, R: GGATCTTGAAGTTGGCCTTG

PCR tag vectors:

pLenti-puro (Addgene 39481; [5]) was modified by PCR to insert a triple Flag tag, followed by the following restriction enzyme sites: NheI, AscI, PacI and BamHI. PCR tags were generated from eYFP by PCR, with six specific pairs of primers and added PacI and BamHI sites, and were inserted into the PacI and BamHI of the modified pLenti-puro. A PacI-XhoI fragment from each of the six pLenti-puro tag vectors was inserted into a modified pLX304 vector (Addgene 25890; [7]), removing the CMV promoter and any polylinker cloning sites. Sequences were verified by Sanger sequencing and are provided as supplementary information. pLX304 was a gift from David Root (Addgene plasmid # 25890; http://n2t.net/addgene: 25890; RRID:Addgene 25890). pLenti-puro was a gift from Ie-Ming Shih (Addgene plasmid # 39481; http://n2t.net/addgene: 39481; RRID:Addgene 39481).

### Other general methods

#### Cell culture and shRNA knock-down

HCT116 and HEK293T cells were cultured in RPMI or DMEM (Invitrogen) with 10% Fetal Bovine Serum (Hyclone). HEK293T cells were transfected with polyethylenimine (PEI) [11], to generate lentiviral supernatants for shRNA knock down with validated Mission lentiviral shRNA vectors (Sigma; TRCN0000020153 - TGIF1). HCT116 cells were transduced with lentiviral vectors under standard conditions using 8μg/ml polybrene (Millipore-Sigma), and 24 hours after infection cells were selected with 0.15μg/ml puromycin for 7 days, when pools were frozen for xenograft generation, and checked for knockdown by western blot, and for the presence of appropriate PCR tags by PCR from gDNA. pLX304 tag vectors were introduced by lentiviral infection as above, and cells were subjected to selection with 3μg/ml blasticidin.

#### Primer design

The six pairs of tag specific primers were based on intron-spanning primer pairs we had previously used for qRT-PCR analysis of gene expression [6], and were selected for having high efficiency. The primer pairs detect cDNA from spliced RNA from ADAMTS5, CCN2, FRMD6, MYT1, MYT1L and FST (for tags 1-6 respectively), but do not efficiently amplify from genomic DNA. All PCR primers were designed using Primer3 (https://primer3.ut.ee/; [12,13]).

#### Orthotopic xenografts

All animal procedures were approved by the Animal Care and Use Committee of the University of Virginia (protocol 4078), which is fully accredited by the AAALAC. Eight week old athymic Foxn1 nude mice (Jackson, 007850) were injected with a total of 1 x 10^6^ cells in the cecal wall at six sites using Hamilton syringe, three injections per side [14]. Mice were anesthetized by IP injection of ketamine/dexmedetomidene mouse mix (80mg/kg and 0.2mg/kg, respectively). Bupivicaine (0.025ml, 0.5%) was injected subcutaneously at the site of incision, 3-5 minutes prior to incision, and the surgical site was wiped with 3 wipes of povidone iodine scrub and 3 wipes of 70% ethyl alcohol prior to starting the incision. The colon was accessed through the incision using a cotton-tipped swab, exteriorized, and held while the cecum was injected with cells. Saline was used to keep the colon moist while exteriorized. Six individual injections of 10μL each were performed on each animal, with a total of 10^6^ cells injected. The colon was placed back in the peritoneal cavity, the abdominal wall incision closed with a single absorbable suture, and the skin closed with wound clips. Atipamezole (2mg/kg, subcutaneous) anesthetic reversal and ketoprofen (4mg/kg, subcutaneous) were immediately administered post wound closure. Mouse health was monitored daily for five days and the wound clips removed at day 7. Tumors were allowed to progress until humane endpoints were reached, or to a maximum of 70 days post tumor inoculation, and then frozen prior to homogenization using Polytron homogenizer at 300 mg/ml in PBS. Criteria monitored for humane endpoints are: extreme decreased mobility, abdominal distension, cachexia, anemia, bloody stool, or losing 20% body weight. Mice were monitored once weekly until tumors were palpable, and then twice weekly. Because we measure the relative proportions of each cell type in any given tumor, and survival was not used as an outcome, mice were euthanized immediately on signs of distress. Two of 24 mice were euthanized due to signs of distress in the week following surgery. The remaining 22 mice were euthanized on initial signs of distress or at a maximum of 70 days. Because this approach measures the relative proportions of each cell type in a given tumor, and survival is not a primary outcome, mice were euthanized immediately on signs of distress.

## Acknowledgements

The authors thank the Research Histology Core and Molmart for assistance, members of the Center for Systems Analysis of Stress-Adapted Cancer Organelles for support and advice, and Kevin Janes for helpful discussion of statistical approaches.

## Supporting information

**S1 Fig. Analysis of qPCR performance.** A) Example of reduced representation mix for Tag 1. Similar mixes for each of the other five tags were also analyzed. Mix “A” contained 10ng of each plasmid in a total of 60ul, in mix “B” and “C”, the amount of one tag plasmid (in this example Tag 1) was reduced to 0.1ng (mix “B”) or 0.01ng (mix “C”). Mix “A” was analyzed by qPCR for all six tags, mix “B” and mix “C” were analyzed by qPCR for the reduced representation tag (Tag 1 in the example), and all mixes were analyzed by qPCR for the common Venus fragment. qPCR was carried out with two amounts of total input (0.1ng and 0.01ng). B) Total input amounts in each mix, total concentration and % of total for the reduced representation tag (in this example Tag 1) is shown, together with the amount (in pg) of the reduced representation plasmid present with either 0.1ng or 0.01ng total input to the qPCR reaction. C) qPCR data testing linearity of reduced representation samples for each of the six tags based on the scheme in panels A and B. Graphs show the log2[input] for the reduced representation tag (in pg) plotted on the x-axis and the delta Ct for [Venus]-[specific Tag] on the y-axis. Blue lines from 0.1ng input, orange lines from 0.01ng input. D) Analysis of the effect of the pre-amplification PCR. From an equal mix of all six tags, 8, 10 or 12 cycles of pre-amplification were performed, followed by column purification. Each pre-amplified sample was analyzed by qPCR for all six tags and the common Venus amplicon. Raw Ct values are plotted against the number of cycles of pre-amplification. Note, that since there is no normalization to a starting sample here, the Cts for each Tag are not expected to be equal, although the Ct for Venus is lower since it is present in all six.

**S1 Table. Analysis of multiple tumors from small numbers of animals.** HT29 cells with either Tag 3 and shControl, or Tag 6 and shACADM were mixed at an equal ratio and used to generate cecal tumors in four mice (summarized bottom left). Data are shown for each tumor (with animal id), as follows: The raw Ct values for Tag 3, Tag 6 and Venus; the ΔCt values for each (calculated as the Ct for the starting mix [s0] – the Ct for each tumor). Note the raw Cts for the S0 sample are shown below those for the tumors; the ΔΔCt values for each Tag (calculated as the ΔCt[Tag] - ΔCt[venus] for each tumor); the proportion of each tag in the tumor. Note the proportions can be calculated either from the ΔCt or the ΔΔCt values, giving the same result. Bottom right shows the p-value calculated by Wilcoxson signed rank, ignoring the fact that multiple tumors come from two of the mice. The p-values calculated by ANOVA (with animal id as a covariate) using the ΔΔCt values are also shown.

**S1 File. Detailed protocol for qPCR analysis of mixed tumors.** A step-by step protocol for qPCR analysis of mixed cell type tumors, metastases, or other competitive growth tests is provided.

## References

1. Voskoglou-Nomikos T, Pater JL, Seymour L. Clinical predictive value of the in vitro cell line, human xenograft, and mouse allograft preclinical cancer models. Clin Cancer Res. 2003;9: 4227–4239.

2. McIntyre RE, Buczacki SJA, Arends MJ, Adams DJ. Mouse models of colorectal cancer as preclinical models. Bioessays. 2015;37: 909–920. doi:10.1002/bies.201500032

3. Lee WY, Hong HK, Ham SK, Kim CI, Cho YB. Comparison of colorectal cancer in differentially established liver metastasis models. Anticancer Res. 2014;34: 3321–8.

4. Cespedes MV, Espina C, Garcia-Cabezas MA, Trias M, Boluda A, Gomez del Pulgar MT, et al. Orthotopic microinjection of human colon cancer cells in nude mice induces tumor foci in all clinically relevant metastatic sites. Am J Pathol. 2007;170: 1077–85. doi:10.2353/ajpath.2007.060773

5. Guan B, Wang TL, Shih Ie M. ARID1A, a factor that promotes formation of SWI/SNF-mediated chromatin remodeling, is a tumor suppressor in gynecologic cancers. Cancer Res. 2011;71: 6718–27. doi:10.1158/0008-5472.CAN-11-1562

6. Melhuish TA, Kowalczyk I, Manukyan A, Zhang Y, Shah A, Abounader R, et al. Myt1 and Myt1l transcription factors limit proliferation in GBM cells by repressing YAP1 expression. Biochim Biophys Acta Gene Regul Mech. 2018;1861: 983–995. doi:10.1016/j.bbagrm.2018.10.005

7. Yang X, Boehm JS, Yang X, Salehi-Ashtiani K, Hao T, Shen Y, et al. A public genome-scale lentiviral expression library of human ORFs. Nat Methods. 2011;8: 659–661. doi:10.1038/nmeth.1638

8. Shah A, Melhuish TA, Fox TE, Frierson HF, Wotton D. TGIF transcription factors repress acetyl CoA metabolic gene expression and promote intestinal tumor growth. Genes Dev. 2019;33: 388–402. doi:10.1101/gad.320127.118

9. Melhuish TA, Adair SJ, Shah A, Bauer TW, Wotton D. Increased expression of a subset of genes within reduced copy number regions across multiple cancer types. bioRxiv: The Preprint Server for Biology; 2026. doi:10.64898/2026.04.10.717791

10. Truett GE, Heeger P, Mynatt RL, Truett AA, Walker JA, Warman ML. Preparation of PCR-quality mouse genomic DNA with hot sodium hydroxide and tris (HotSHOT). Biotechniques. 2000;29: 52, 54.

11. Boussif O, Lezoualc’h F, Zanta MA, Mergny MD, Scherman D, Demeneix B, et al. A versatile vector for gene and oligonucleotide transfer into cells in culture and in vivo: polyethylenimine. Proc Natl Acad Sci USA. 1995;92: 7297–7301. doi:10.1073/pnas.92.16.7297

12. Koressaar T, Remm M. Enhancements and modifications of primer design program Primer3. Bioinformatics. 2007;23: 1289–1291. doi:10.1093/bioinformatics/btm091

13. Untergasser A, Cutcutache I, Koressaar T, Ye J, Faircloth BC, Remm M, et al. Primer3--new capabilities and interfaces. Nucleic Acids Res. 2012;40: e115. doi:10.1093/nar/gks596

14. Chen GT, Tifrea DF, Murad R, Habowski AN, Lyou Y, Duong MR, et al. Disruption of β-Catenin-Dependent Wnt Signaling in Colon Cancer Cells Remodels the Microenvironment to Promote Tumor Invasion. Mol Cancer Res. 2022;20: 468–484. doi:10.1158/1541-7786.MCR-21-0349

