## Supplemental Figure for "A simple method for analyzing competitive growth of multiple cell types in xenograft tumors"

A

One equal mix, and two mixes with reduced representation of each tag (13 total).

| Tag | 1 | 2 | 3 | 4 | 5 | 6 |
|---|---|---|---|---|---|---|
| Mix A | 10 | 10 | 10 | 10 | 10 | 10 |
| Mix B | 0.1 | 10 | 10 | 10 | 10 | 10 |
| Mix C | 0.01 | 10 | 10 | 10 | 10 | 10 |

qPCR from each for reduced representation Tag, Venus, and all six Tags from equal mix, with two input amounts (0.1ng and 0.01ng total).

B

| Mix | A | B | C |
|---|---|---|---|
| Input Tag 1 (ng) | 10 | 0.1 | 0.01 |
| Input Tags 2-6 (ng ea) | 10 | 10 | 10 |
| Total input (ng) | 60 | 50.1 | 50.01 |
| Concentration (ng/ul) | 10 | 8.35 | 8.335 |
| Tag 1 (% total) | 16.67 | 0.1996 | 0.02000 |
| pg Tag 1 (0.1ng PCR) | 16.7 | 0.167 | 0.0167 |
| pg Tag 1 (0.01ng PCR) | 1.67 | 0.0167 | 0.00167 |

C

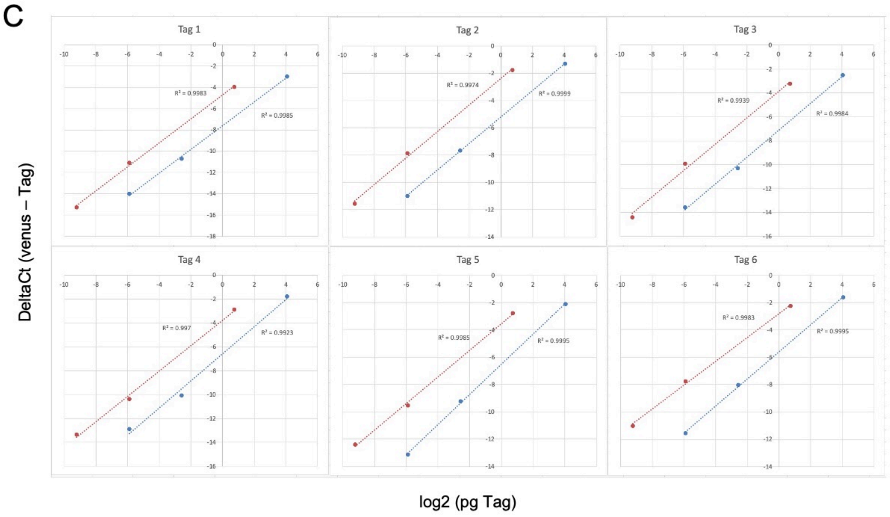


D

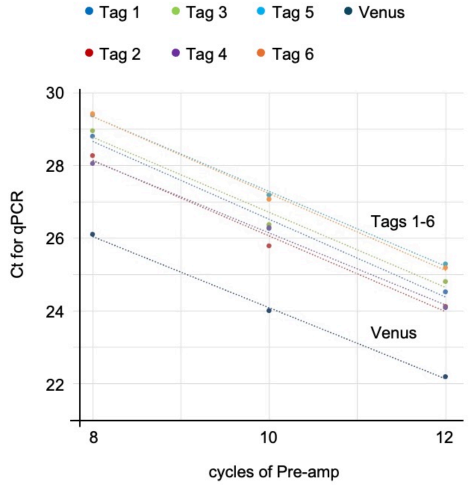
