## Supplemental Table for "A simple method for analyzing competitive growth of multiple cell types in xenograft tumors"

| | | Ct | | | dCt (S0-tumor) | | | ddCt (tag-venus) | | | |
|---|---|---|---|---|---|---|---|---|---|---|---|
| animal_id | tumor_id | Tag 3 | Tag 6 | Venus | Tag 3 | Tag 6 | Venus | Tag 3 | Tag 6 | prop T3 | prop T6 |
| 1 | 1 | 30.660 | 30.020 | 28.370 | -6.207 | -5.183 | -5.330 | -0.877 | 0.147 | 0.3297 | 0.6703 |
| 2 | 2 | 29.220 | 28.690 | 27.220 | -4.767 | -3.853 | -4.180 | -0.587 | 0.327 | 0.3468 | 0.6532 |
| 2 | 3 | 29.630 | 29.030 | 27.460 | -5.177 | -4.193 | -4.420 | -0.757 | 0.227 | 0.3359 | 0.6641 |
| 2 | 4 | 26.590 | 26.110 | 24.730 | -2.137 | -1.273 | -1.690 | -0.447 | 0.417 | 0.3547 | 0.6453 |
| 2 | 5 | 24.910 | 24.020 | 23.060 | -0.457 | 0.817 | -0.020 | -0.437 | 0.837 | 0.2926 | 0.7074 |
| 3 | 6 | 25.850 | 25.180 | 23.760 | -1.397 | -0.343 | -0.720 | -0.677 | 0.377 | 0.3252 | 0.6748 |
| 3 | 7 | 25.860 | 25.420 | 23.970 | -1.407 | -0.583 | -0.930 | -0.477 | 0.347 | 0.3611 | 0.6389 |
| 3 | 8 | 31.820 | 31.720 | 29.450 | -7.367 | -6.883 | -6.410 | -0.957 | -0.473 | 0.4170 | 0.5830 |
| 4 | 9 | 26.240 | 26.780 | 24.450 | -1.787 | -1.943 | -1.410 | -0.377 | -0.533 | 0.5271 | 0.4729 |

| | | | |
|---|---|---|---|
| S0 | 24.453 | 24.837 | 23.040 |

HT29 cecal tumors
1:1 mix of Tag 3 cells (shCtrl) and Tag 6 cells (shACADM)
Four mice, with a total of 9 tumors

| Mouse # | # tumors |
|---|---|
| 1 | 1 |
| 2 | 4 |
| 3 | 3 |
| 4 | 1 |

Wilcoxson signed rank, using proportions
p-value 0.007812

ANOVA using ddCt values
p-value by animal 0.331
p-value by Tag 0.000152
